# Interdependent photo- and chemosensory systems regulate larval settlement in a marine sponge

**DOI:** 10.1101/519512

**Authors:** Tahsha E. Say, Sandie M. Degnan

## Abstract

Marine pelagic larvae from throughout the animal kingdom use a hierarchy of environmental cues to identify a suitable benthic habitat on which to settle and metamorphose into the reproductive phase of the life cycle. The majority of larvae are induced to settle by biochemical cues (1) and many species have long been known to preferentially settle in the dark (2). Combined, these data suggest that larval responses to light and biochemical cues may be linked, but this is yet to be explored at the molecular level. Here, we track vertical position of larvae of the sponge *Amphimedon queenslandica* to show that they descend to the benthos at twilight, by which time they are competent to respond to biochemical cues (3), consistent with them naturally settling in the dark. We then conduct larval settlement assays under three different light regimes (natural day-night, constant dark or constant light), and use transcriptomics on individual larvae to identify candidate molecular pathways underlying the different settlement responses that we observe. We find that constant light prevents larval settlement in response to biochemical cues, likely via actively repressing chemostransduction; this is consistent with the sustained upregulation of a photosensory cryptochrome and two putative inactivators of G-protein signalling in the constant light only. We hypothesise that photo- and chemosensory systems may be hierarchically integrated into ontogeny to regulate larval settlement via nitric oxide (NO) and cyclic guanosine monophosphate (cGMP) signalling in this sponge that belongs to one of the earliest branching of the extant animal lineages.

**Significance statement:** In the ocean, successful recruitment of pelagic larvae into reproductive adult populations enables the survival and connectivity of benthic communities. The majority of invertebrate larvae are induced to settle by biochemical cues, and multiple species preferentially settle in the dark. Here, we explore, for the first time, interactions between light and biochemical cues at behavioural and molecular levels during larval ontogeny in a sponge. We find that light perturbs ontogenetic changes in gene expression and prevents settlement in response to biochemical cues, demonstrating strong interdependencies between photo- and chemosensory systems. Sponges are one of the earliest-branching of the extant animal phyletic lineages, and a valuable comparative model for understanding the origin and evolution of the pelago-benthic life cycle.

## Introduction

Recruitment of marine invertebrate larvae into reproductive adult populations is often contingent on the irrevocable pelago-benthic transition. To achieve this, larvae must descend to the bottom of the water column and identify a suitable benthic substrate on which to settle; they do so by using a hierarchy of physical and biological environmental cues (reviewed in 4). The physical cue of light often influences larval swimming orientation and settlement (2, 5), but larvae of most species will not settle until they encounter an appropriate biological cue, typically a biochemical cue associated with coralline algae (1). To respond to biochemical settlement cues, larvae must first develop a functional chemosensory system, defined as the acquisition of competence (1). Competence has long been hypothesised to be acquired upon the accumulation of a threshold concentration of chemosensory G-protein coupled receptors (GPCRs) (6). More recently, it has also been suggested that the acquisisiton of competence can be accelerated by an environmental physical cue (7, 8), but the underlying molecular mechanisms remain to be explored. Here, we combine behavioural and molecular assays to explore, for the first time, how responses to physical and biochemical cues are integrated during larval ontogeny, using a representative of an early-branching animal lineage, the demosponge *Amphimedon queenslandica*.

Competent *A. queenslandica* larvae do not settle unless they encounter an appropriate biochemical cue (9), such as the articulated coralline algae *Amphiroa fragilissima* (10). *A. queenslandica* larvae emerge from maternal brood chambers throughout the early to mid afternoon (11, 12). The majority are competent to respond to biochemical cues by 4-6 hours post emergence (3), which coincides with twilight, suggesting that these larvae naturally settle in the dark. Indeed, sponge larvae are known to prefer shaded surfaces for settlement (13) and this is consistent with *A. queenslandica* larvae being negatively phototactic immediately post-emergence (14, but see 3). The phototaxis is regulated by a blue-light flavoprotein, a cryptochrome, which is localised to cells of the posterior photosensory organ in precompetent larvae, prior to their emergence from the adult (12, 15). These observations raise the hypothesise that two interdependent sensory systems – a photosensory cryptochrome and a chemosensory system – orchestrate the pelago-benthic transition in this species.

Here, we test the interactions between photo- and chemosensory systems at both behavioural and transcriptomic levels in *A. queenslandica* larvae. First, we disentangle the effect of light (a physical cue) and *A. fragilissima* (a biochemical cue) on larval behaviour using vertical swimming and settlement assays. Second, we examine the molecular interdependencies between photo- and chemosensory systems by measuring global transcriptional changes in individual precompetent and competent sponge larvae that were reared under one of three experimental light regimes (natural day-night, constant dark or constant light). In doing so, we generate the first molecular data to explore how physical and biochemical cues are hierarchically integrated into larval ontogeny to regulate the pelago-benthic transition. We pay particular attention to the expression of the widely-conserved chemosensory GPCRs and the nitric oxide (NO) signalling system because of their hypothesised role in the settlement of multiple marine invertebrates, including this sponge (16–21). Sponges have existed for more than 700 million years (22) and are one of the earliest branching of the extant animal phyletic lineages (23, 24) making them a valuable comparative model for understanding conserved molecular mechanisms orchestrating larval settlement throughout the animal kingdom.

## Results

### Larval vertical position in the water column is influenced by ontogenetics interacting with a hierarchy of environmental cues

#### Negative phototaxis directs larval distribution in the water column before twilight

To determine how light influences the vertical position of larvae through time, we commenced vertical swimming assays within one hour post emergence from adult sponges beginning at 12 pm, 1 pm and 2 pm. Larvae were given a choice of swimming within or outside a shaded region that was positioned at either the top, middle or bottom of the water column. On average, 94%, 82% and 89% of *A. queenslandica* larvae were located within the dark-shaded regions of the water column at 2, 3 and 4 pm, respectively (Fig. 1*A*). At twilight (6:30 pm), larvae (of mixed ages and regardless of shade location) had begun to migrate down towards the bottom of the water column (Fig. 1*A*). Only 41% and 23% of larvae were positioned in the shaded top and middle regions of the column at 7 pm, respectively, and this later decreased to 15% and 12% by 10 pm; at 11 pm, no larvae were present in the top- or middle- shaded regions (Fig. 1*A*). In contrast, larvae (74%) were concentrated in the shaded-bottom of the column at 7 pm, and this further increased to 89% and 95% at 10 and 11 pm, respectively (Fig. 1*A*). There was a significant interaction between time and the mean proportion of larvae occupying the shaded-regions of the water column, consistent with a change in larval behaviour upon the removal of darkness as a directional cue at twilight (repeated-measures ANOVA, ε = 0.399, F_3, 38_ = 22, P<0.001). Bonferroni pairwise comparisons revealed a significant difference in the proportion of larvae occupying the shaded bottom compared to the shaded top (mean difference = 1.889, P<0.001) and shaded middle (mean difference = 2.733, P<0.001) of the column.

**Fig. 1.**
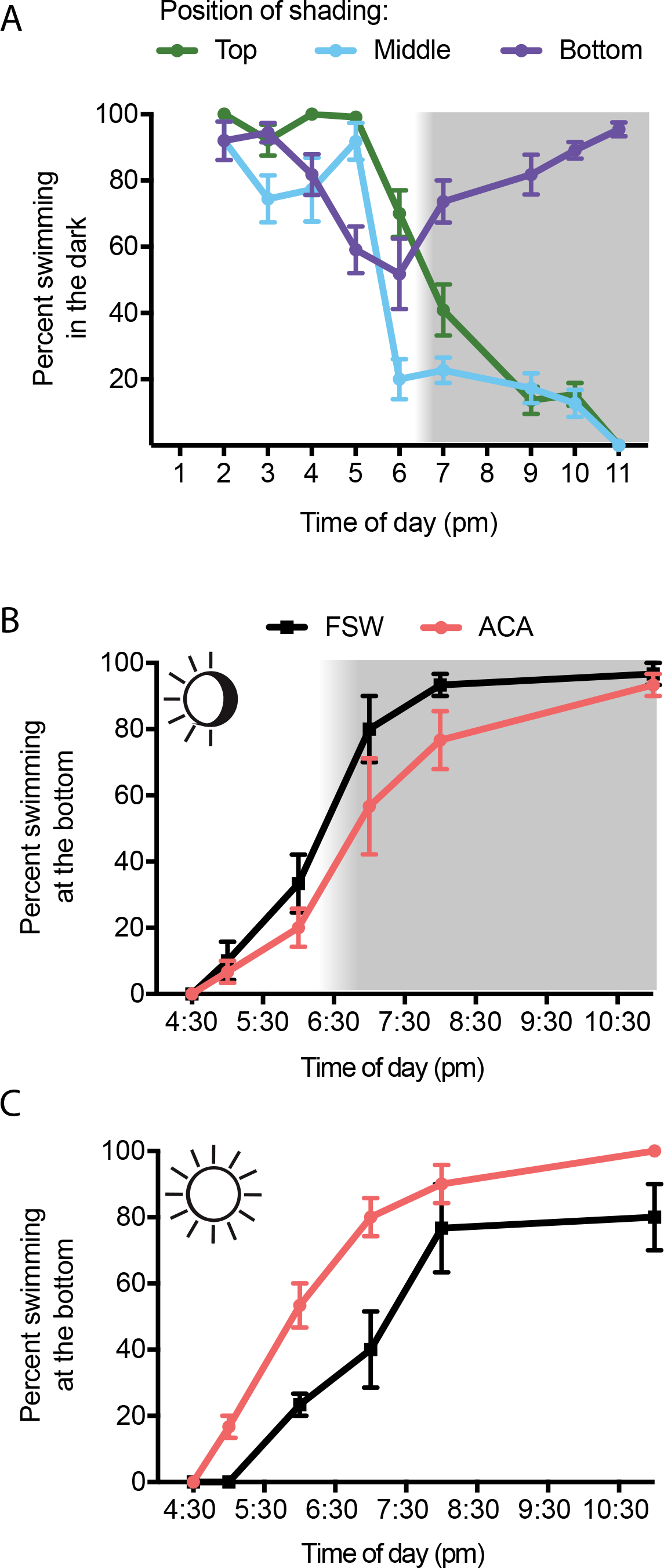
Larvae descend to the bottom of the water column at twilight. (A) Percent (mean ± SE, n = 5, 9 and 11 columns at 2, 3 and 4-11 pm, respectively) of *Amphimedon queenslandica* larvae in the dark-shaded regions positioned at either the top, middle or bottom third of the water column containing 0.22 μm filtered sea water (FSW) and 10 larvae. (B) Mean (±SE, n = 3) percentage of larvae in the bottom of the water column in natural day-night and (C) constant light, in the presence (pink) of an effective biochemical settlement cue, the articulated coralline algae (ACA) *Amphiroa fragilissima* or the FSW negative control (black), only. Shading represents hours of darkness following sunset.

#### Larvae descend to the bottom of the water column at twilight

The mean proportion of *Amphimedon queenslandica* larvae at the bottom of vertical swimming assays significantly differed over time (repeated-measures ANOVA, ε = 0.494, F_2, 20_ = 145, P<0.001), and there was no significant interaction with biochemical settlement cue and/or light regime. Specifically, larvae migrated down towards the bottom around twilight (6:20 pm), irrespective of whether a biochemical settlement cue – the articulated coralline algae (ACA) *Amphiroa fragilissima* – was present or absent (Fig. 1 *B* and *C*). Under natural day-night, the percentage of larvae occupying the bottom of the water column increased from 33% to 93% from 6 pm to 8 pm in the absence of a biochemical cue (Fig. 1*B*). Similarly, in the presence of a biochemical cue, 20% to 77% of larvae were swimming at the bottom at 6 pm and 8 pm, respectively (Fig. 1*B*). This downward migration corresponding with twilight was maintained even in a constant light regime, with 90% and 77% of larvae located at the bottom at 8 pm in the presence and absence of a settlement cue, respectively (Fig. 1*C*). Post-hoc tests using the Bonferroni correction revealed that the mean proportion of larvae at the bottom of the water column, across all experimental light regimes, significantly increased from 4:30 pm until 40 minutes after twilight (P< 0.05*, P<0.01**, P<0.005***, mean difference ±SE: 4:30-5 pm*, 0.833±0.186; 5-6 pm***, 2.4176±220; 6-7 pm**, 3.167±0.527).

### Larvae do not settle in response to biochemical cues when maintained in constant light

In the sponge *Amphimedon queenslandica*, the timing and percentage of larval settlement in response to the ACA *Amphiroa fragilissima* varied under the three different light regimes that were tested (natural day-night, constant light and constant dark). In the natural day-night regime, 100% of larvae were swimming at 4 hpe, three hours before twilight (7 hpe; Fig. 2*A*). However, at twilight (7 hpe), percent swimming decreased to 74%, indicating that 26% of larvae had settled in response to the ACA (Fig. 2*A*). Percent swimming further decreased to <50% within just two hours of twilight (at 9 hpe) and just 20% swimming at 13 hpe (Fig. 2*A*). At 6 hpe (one hour before twilight), irradiance levels in natural day-night measured 2.54 ± 0.26 μmol photons · m^−2^ · s^−1^, still higher than the constant dark level of 0 μmol photons · m^−2^ · s^−1^. In constant dark, larvae initiated settlement within just 1 hpe and high settlement (<50% swimming) was observed by 7 hpe; only 38% remained swimming at 13 hpe (Fig. 2*B*). In contrast, in constant light, only 2% of larvae had settled in response to the ACA at the conclusion of the experiment at 13 hpe (Fig. 2*C*). The artificial light intensity in the constant light regime averaged ~13.56 ± 0.24 μmol photons · m^−2^ · s^−1^ which corresponds to irradiance levels at 5 hpe, 2 hours before twilight (at 7 hpe) under a natural day-night regime (~13.25 ± 0.29 μmol photons · m^−2^ · s^−1^). To determine if the settlement behaviour observed in constant light could be altered by transitioning larvae into the dark, a subset of larvae were transferred from constant light into darkness at 10 hpe (see Methods). Importantly, after just one hour since transfer from constant light into darkness, only 61% of larvae remained swimming and this further decreased to 38% within another two hours (Fig. 2*C*); this was significantly different to settlement of larvae maintained in constant light (logrank test; χ^2^ = 11.2 on 1 DF, P < 0.005).

**Fig. 2.**
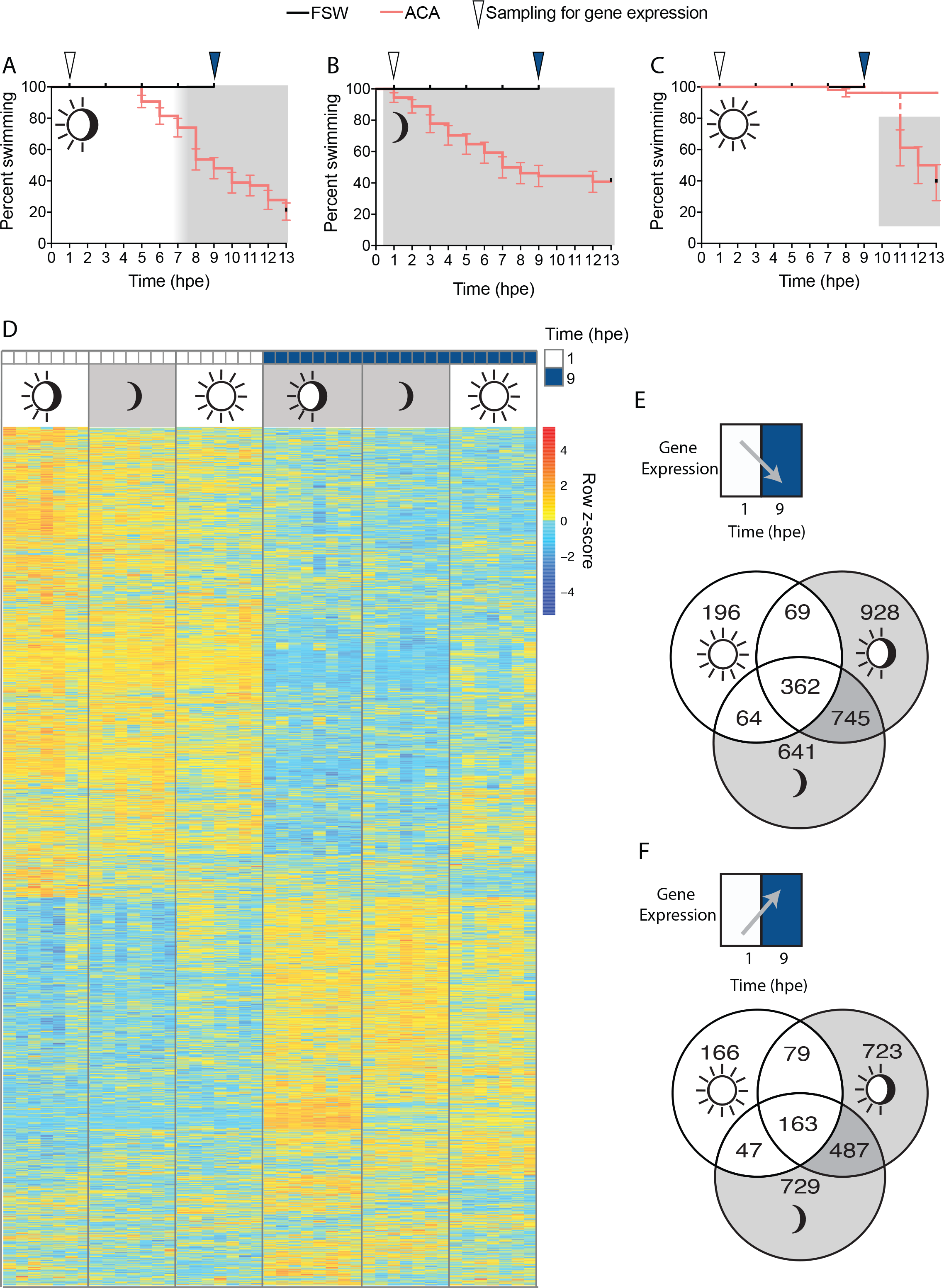
Constant light retards larval settlement. Percent (± SE) swimming *A. queenslandica* larvae under (A) natural day-night, (B) constant dark, and (C) constant light regimes (n = 54 for each). Larvae were distributed into settlement assays within 0.5 hours post emergence into daylight (hpe) and a single larva was exposed to either the articulated coralline algae (ACA) *Amphiroa fragilissima* as the settlement cue (pink) or 0.22 μm filtered sea water (FSW) negative controls (black). Grey shading represents hours of darkness. In (C) larvae (n = 18 individual larvae) were transferred from constant light into darkness at 10 hpe. Arrows represent the times chosen for gene expression analysis at 1 and 9 hpe (white and blue, respectively). (D) Normalised gene expression of all 5799 significantly differentially expressed genes in response to light regime and/ or time (hpe) in individual larvae (n = 7-8) identified using the LRT analysis in DESeq2. This heatmap illustrates z-scores representing standard deviations from the row mean thus allowing relative comparisons among individual larvae for each gene. Red and dark blue indicates highest and lowest levels of gene expression, respectively. The total numbers of genes that significantly (E) decreased and (F) increased in expression between 1 and 9 hpe were independently assessed for each light regime using the Wald test in DESeq2. All DEGs were identified at the 0.05 significance level.

Larval settlement in response to the ACA significantly increased with larval age (GLM; P < 0.005; Fig. 2*B*); all time points except 2 hpe significantly differed from 1 hpe. However, regardless of age, larvae were significantly more likely to respond to the ACA in the natural day-night and constant dark regimes compared to constant light (GLM; z = 11.31, P <0.005 and z = 12.82, P < 0.005, respectively). Larvae were also significantly more likely to settle in constant dark compared to a natural day-night regime (z = 3.58, P < 0.0005).

In the filtered sea water (FSW) negative controls, no settlement was observed at any time in any light regime by 9 hpe (Fig. 2 *A*–*C*), at which time the last of the larvae were sampled for gene expression analyses (see Methods). Thus there was significantly lower settlement in the FSW negative controls compared to when larvae were presented with ACA (logrank test; χ^2^ = 26 on 1 DF, P < 0.005).

### Although ontogeny drives the majority of the changes in larval transcriptomes, light influences the expression of some critical genes

To understand the molecular interdependencies between photo- and chemosensory systems in *A. queenslandica* larvae, we analysed genome-wide gene expression in individual precompetent and competent larvae from each of the three different experimental light regimes. On average, 77% of the reads sequenced using CEL-Seq2 (25), an RNA-Seq approach, successfully mapped to the genome (Dataset S1). Raw counts are available as a supplementary dataset (Dataset S2). Principal component analyses (26) revealed that transcriptional changes during larval ontogeny were more pronounced after a light-to-dark transition at twilight, compared to larvae that were maintained in constant light and constant dark, accounting for 15% of explained variation in larval transcriptomes (Fig. S1 *A*–*D*).

To explore how gene expression is influenced by larval aging and/ or light regime we performed Likelihood Ratio Test (LRT) statistics in DESeq2 (27). Of the 47,411 protein coding and long non-coding RNAs (lncRNAs) in *A. queenslandica* (28, 29), 26,450 were expressed in this CEL-seq2 experiment after filtering. Of these, only 8% (5598 gene models and 139 lncRNAs) were significantly differentially expressed among any light regime and/or time (hpe), including any interactions (Fig. 2*D*, Dataset S3). Ontogeny drove most of the changes in gene expression, as indicated by the distribution of the significantly differentially expressed genes (DEGs) into two major K-Means clusters according to whether their highest expression is at 1 or 9 hpe (Fig. 2*D*). However, at 9 hpe, light regime also significantly affected gene expression; specifically, larvae reared in natural day-night or in constant dark had more similar gene expression profiles to each other compared to those reared in constant light (Fig. 2*D*). This is consistent with the former also having more similar settlement rates to each other (Fig. 2 *A–C*).

To identify changes in gene expression between 1 and 9 hpe, independently for each light regime, we utilised the Wald test in DESeq2 (27). The DEG lists were compared across light regimes using Venny 2.0 (30) and we did this separately for DEGs with a higher expression at either 1 hpe (negative fold change) or 9 hpe (positive fold change; Fig. 2 *E* and *D*). We found that 362 DEGs were more highly expressed at 1 hpe, and 163 DEGs were more highly expressed at 9 hpe irrespective of light regime (Fig. 2 *E* and *F*). Importantly, we found also that constant light retards transcriptional changes through ontogeny (Fig. 2 *E* and *F*). In constant light, only 196 DEGs decreased between 1 and 9 hpe, and only 166 DEGs increased (Fig. 2 *E* and *F*). However, after larvae experienced a light-to-dark transition that occurred either at twilight (in the natural day-night regime) or immediately upon emergence (in the constant dark regime), 745 DEGs decreased and 487 DEGs increased in expression over this same time period (Fig. 2 *E* and *F*, dark shading). The difference in the timing of the light-to-dark transition itself also influenced transcriptional changes. A total of 928 and 641 unique DEGs were downregulated at 1 hpe in the natural day-night and constant dark regimes, respectively (Fig. 2*E*); and 723 and 729 DEGs were upregulated at 9 hpe in the natural day-night and constant dark regimes, respectively (Fig. 2*F*).

#### Light perturbs ontogenetic changes in gene expression

To identify the genes that are most strongly influenced by light regime and larval age, we used the supervised sparse partial least squares discriminant analysis (sPLS-DA; 31) which identified 178 gene models and 2 lncRNAs that alone explain 9% of the variation (Fig. 3*A*, C1). We see no effect of light on the expression of these genes at 1 hpe (Fig. 3*A*). This is consistent with larvae having only spent half an hour in their respective light regimes, because all larvae emerged into natural daylight before being distributed equally into one of three experimental light regimes by 0.5 hpe. Importantly constant light appears to retard ontogenetic changes in gene expression from 1 hpe to 9 hpe (Fig. 3*A*), and this is consistent with the majority of larvae being unable to respond to biochemical settlement cues in the light (Fig. 2*C*). Larvae aged 9 hpe in the natural day-night and constant dark regimes show similar expression profiles to each other (Fig. 3*A*, C1). We find that just 20 genes, explaining 3% of the variability, show differences in expression depending on the timing of the light-to-dark transition; this transition occurred at either 0.5 hpe when larvae were introduced into the constant dark regime, or at twilight in the natural day-night regime (Fig. 3*A*, C2). However, because both of these light regimes involved a light-to-dark transition and yielded high rates of larval settlement (Fig. 2 *A* and *B*), all subsequent analyses were performed on the genes identified on the first component only. We found that 78% of these genes (i.e. 141 gene models and lncRNAs) were more highly expressed at 1 hpe, compared to 9 hpe, as indicated by the two major K-means clusters of the hierarchically-clustered heat map (Fig. 3*B*). All of the genes identified by the sPLS-DA were also previously identified as being significantly differentially expressed by the LRT statistics (Dataset S4), including the photoreceptor gene *cryptochrome AqCry2* that was the most significantly DEG over ontogeny in both the natural day-night and the constant dark regimes (Fig. 4*A*). Complete DEG lists with Blast2GO, InterPro and KEGG annotations are listed in (Dataset S5), and with Pfam annotations in (Dataset S6).

**Fig. 3.**
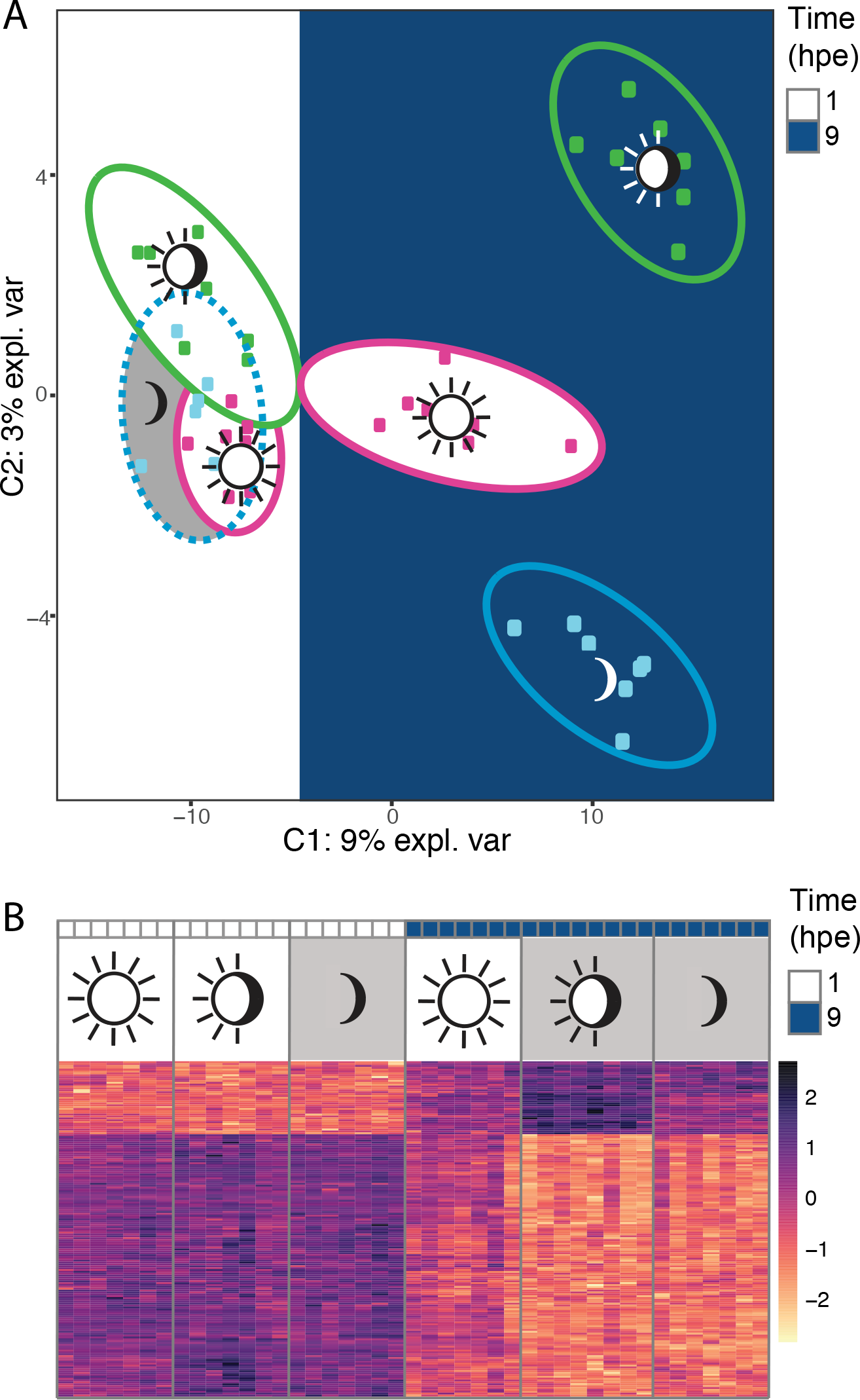
A supervised analysis reveals that light retards changes in gene expression during ontogeny in the sponge *Amphimedon queenslandica* larvae. (A) Supervised sparse partial least square discriminant analysis (sPLS-DA) identified 180 gene models and long non-coding RNAs on C1, and 20 genes on C2, that alone explain 9% and 3% of explained variability. Precompetent and competent larvae (n = 7-8) were aged 1 and 9 hours post emergence (hpe) at the time of sequencing, indicated by white and blue shading, respectively; larvae were reared under one of three different light regimes (constant light, natural day-night and constant dark). Ellipses indicate 95% confidence intervals. (B) Expression of the 180 genes identified on C1 visualised on a hierarchically clustered heatmap using variance stabilising transformed (vst) counts. This heatmap illustrates z-scores representing standard deviations from the row mean thus allowing relative comparisons among individual larvae for each gene. Blue and yellow indicates highest and lowest levels of gene expression, respectively.

**Fig. 4.**
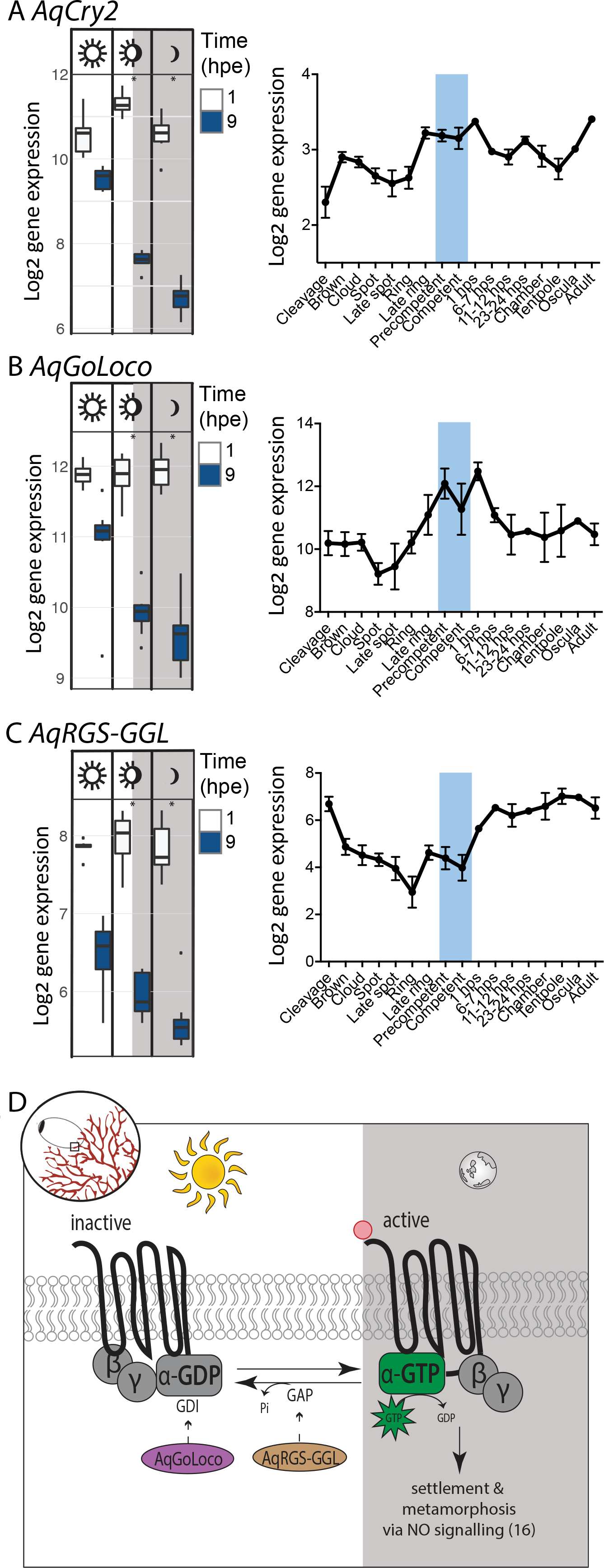
Larval settlement is orchestrated by interdependent photo- and chemo- sensory systems in the sponge *Amphimedon queenslandica*. Expression of candidate genes involved in *A. queenslandica* larval settlement, (A) the photoreceptor *AqCry2* and two repressors of G-protein signalling, (B) *AqGoLoco* and (C) *AqRGS-GGL*. Box plots show expression in larvae (n = 7-8) aged 1 and 9 hours post emergence (hpe) that were subjected to one of three different light regimes (constant light, natural day-night and constant dark). Boxes indicate the interquartile range (IQR) around the median of log_2_ normalised expression and whiskers extend to 1.5 times the IQR. Asterisks represent genes that are significantly differentially expressed over a threshold of 1 (on the log_2_ scale) between 1 and 9 hpe. Log_2_ developmental expression profiles (mean ± SE) throughout the entire *A. queenslandica* life cycle were based on 82 CEL-Seq samples (41, 42; NCBI GEO GSE54364) comprising 17 stages of development. Cleavage to late ring are embryonic stages; precompetent and competent are larvae aged 0-7 hpe and 6-50 hpe, respectively (blue shading); 1 h post-settlement (1 hps) to tentpole are metamorphic stages; juvenile; adult. (D) Standard model of guanine nucleotide cycle of GPCRs adapted from (89) and contextualised during larval settlement in the sponge *A. queenslandica*. Conformational change between active and inactive states of GPCRs is initiated via binding of an exogenous ligand to a G-protein coupled receptor and in this case represents a larva encountering a biological settlement cue (illustrated in the top left hand corner). GoLoco functions as a guanine dissociation inhibitor (GDI) for Gα subunits, inhibiting the exchange of GDP for GTP and thus blocking Gα signal transduction (34). Regulators of G-protein signalling (RGS) inactivate Gα signalling via binding to active GTP-bound Gα subunits and accelerating their intrinsic GTPase activity to return to a GTP-bound state and terminating the transduced signal allowing them to reassociate with the receptor for subsequent rounds of signalling (35). This general scheme is based on patterns gene expression from individual larval transcriptomes in *A. queenslandica*. Sun and moon symbols are courtesy of the Integration and Application Network (ian.umces.edu/symbols/).

To characterise trends in biological function, we identified functional protein domains that were enriched in the genes identified by the sPLS-DA (Fig. S2). The GoLoco motif (32), also known as the G protein regulatory (GPR) motif (33) was the most significantly enriched Pfam domain (enrichment score=134.4, P < 0.0005; Fig. S2). In particular, the sPLS-DA list included a single gene containing four GoLoco motifs (Aqu2.1.22573_001; XP_003389018.1), hereafter named *AqGoLoco*. The GoLoco motif functions as a guanine dissociation inhibitor (GDI) for Gα subunits, inhibiting the exchange of GDP for GTP and thus blocking G protein signal transduction; it commonly occurs in tandem repeats (34) (Fig. 4*A*). Further investigation of the sPLS-DA list revealed a gene (Aqu2.1.36017_001; XP_019850380.1) encoding a Regulator of G-protein signalling (RGS) domain that is hallmark to another family of proteins that also negatively regulate G-protein signalling (35). The RGS domain controls the timely inactivation of G-protein signalling via binding to active GTP-bound Gα subunits and accelerating their intrinsic GTPase activity, returning them to an inactive state; this terminates signal transduction and allows the Gα subunits to reassociate with the receptor for subsequent rounds of signalling (35) (Fig. 4*D*). This *A. queenslandica RGS* gene (*AqRGS-GGL*) also contains a DEP domain (36) that may function in membrane localization (37), and a G-protein gamma (γ)-like (GGL) domain that can interact with the G-protein β subunit (38) to inhibit association with the conventional G-protein γ subunit, thereby preventing heterotrimer formation (39).

#### Many components of the NO-cGMP pathway play a role during larval ontogeny

We conducted a KEGG pathway analysis on all genes that were expressed in this CEL-Seq2 dataset, after filtering, to identify all components of the NO-cGMP pathway that are expressed during larval ontogeny (Fig. S4). Nitric oxide synthase *AqNOS* (Aqu2.1.39814_001; XP_011410183) was consistently expressed in the top 90th percentile of genes in these transcriptomes, irrespective of light regime, supporting an important role for nitric oxide (NO) during larval life (Fig. S3). Notably, key components of the cGMP pathway were significantly down-regulated over ontogeny, except in the light; this included a membrane-bound atrial natriuretic peptide (ANP)-activated particulate guanylyl cyclase (ANPR-A), and a sarcoplasmic/ endoplasmic reticulum Ca^2+^ ATPase (SERCA), a plasma membrane Ca^2+^ ATPase (PMCA), and other genes regulating Ca^2+^-dependent cellular functions (Fig. S4). In this sponge, GPCRs are hypothesised to activate NO-cGMP signalling (16). Of the 125 *Rhodopsin* and 32 *Glutamate GPCRs* in *A. queenslandica* (40)(Dataset S7), 70% (88 gene models) and 87% (28 gene models) were expressed in our filtered dataset, respectively (Fig. S5). A total of 18 *Rhodopsin GPCRs* and 12 *Glutamate GPCRs* were differentially expressed between 1 hpe and 9 hpe larvae, and the majority were more highly expressed at 1 hpe; this suggests that *GPCRs* are already active in larvae that have recently emerged from the adult (Fig. S6 *A* and *B*). Importantly, 15 *Rhodopsin* and 9 *Glutamate GPCRs* only showed significant changes in expression after a light-to-dark transition (Fig. S6); two of these 15 *Rhodopsin GPCRs* were also identified in the sPLS-DA list of 180 genes (representing 0.04% of the sponge genome) that were most strongly influenced by light (Dataset S5).

#### *Developmental expression profiles throughout the entire* Amphimedon queenslandica *life cycle are consistent with important roles during larval life*

Using existing developmental transcriptomes that cover the entire *A. queenslandica* life cycle (41, 42; NCBI GEO GSE54364) we determined the expression profiles of the photosensory cryptochrome, *AqCry2*, and two putative negative-regulators of chemosensory G-protein signalling, *AqGoLoco* and *AqRGS-GGL*. The expression of *AqCry2* and *AqGoLoco* starts to increase prior to larval emergence from the adult and this is consistent with an important role for both during larval life (Fig. 4 *A* and *B*). The higher expression of *AqRGS-GGL* and *AqGoLoco* in precompetent larvae (Fig. 4 *B* and *C*), which are refractory to settlement cues, is also consistent with their predicted role in negatively regulating G-protein signalling.

## Discussion

### Larvae descend to the bottom of the water column at twilight, as they near competence

*Amphimedon queenslandica* larvae use a hierarchy of environmental cues to mediate their position in the water column. Prior to sunset, larval swimming direction is largely mediated by the physical cue of light (Fig. 1*A*). This is consistent with *A. queenslandica* larvae being negatively phototactic upon emergence from the adult, with the photosensory pigment ring that is comprised of long posterior cilia and associated cryptochrome-expressing cells controlling this behaviour (14, 15). Interestingly, not *all* larvae in our study were strongly negatively phototactic; ~30% of larvae were not attracted to a shaded-region from 1-3 pm unless it was positioned at the top of the vertical column (Fig. 1*A*). Indeed, it has been previously shown that at least 10% of newly emerged *A. queenslandica* larvae immediately swam up towards the top of the water column (3) and our results here indicate that this is independent of phototaxis. This upward swimming may be a response to the geomagnetic field or the partial pressure gradient of dissolved gases (43). This variability in larval phototactic responses in *A. queenslandica* can have important implications for their dispersal capacity, because larvae that are strongly negatively phototactic and swim towards the benthos are less likely to disperse far away from natal habitats (2).

At twilight, the majority of *A. queenslandica* larvae descend to the sea floor (Fig. 1*B*). This uniform downward migration of larvae, irrespective of age, is not likely the result of a change in larval density due to a depletion of nutritional reserves, because larval age varied by up to 3 hours, or to a change in spicule number, because this remains relatively constant though *A. queenslandica* larval development (44). We observed this descent to the bottom at twilight in all of our experimental light regimes, including constant light (Fig. 1*C*), suggesting that entrained circadian rhythmicity may be governing this behaviour. The temporal and spatial expression of molecular homologs of core circadian genes, including a blue-light flavoprotein, the *cryptochrome (AqCry2),* provides direct support for the role of circadian rhythms in larval development in this sponge (12, 15). Indeed, *cryptochromes* have light-dependent and/or diurnal expression patterns that are consistent with a circadian role in the cnidarians *Nematostella vectensis* (45), *Acropora millepora* (46, 47) and *Favia fragum* (48), as well as in the sponge *Suberites domuncula* (49). More recently, it has been suggested that cryptochromes may also function as light-sensitive chemical-based magnetoreceptors (reviewed in 50), which could explain the uniform downward migration of *A. queenslandica* larvae at twilight. Regardless of the underlying cellular basis, we demonstrate that larvae of this sponge descend to the sea floor as they near competence (3) as has been shown for other subtidal marine invertebrates (51, 52).

### *Amphimedon queenslandica* larvae do not settle if maintained in constant light

After descending to the sea floor, *A. queenslandica* larvae must detect an appropriate biochemical cue that induces settlement (9). Here, we demonstrate that they cannot respond to a highly inductive biochemical cue when maintained in constant light, indicating that the photosensory system overrides larval responses to biochemical cues (Fig. 2*C*). This interpretation is further supported by the observation that larvae transferred from constant light into darkness at 10 hours post emergence (hpe) were 8.9 times more likely to settle in response to a biochemical settlement cue compared to larvae that remained in constant light, and 61% did so within just one hour of darkness (Fig. 2*C*, shading). This rapid pelago-benthic transition in response to environmental cues is consistent with the ‘need for speed’ hypothesis suggested to maximize larval and post-larval performance (1), and highlights the importance of two interdependent sensory systems for orchestrating larval settlement in this sponge. In the next sections, we discuss the first molecular evidence of this interdependency between photo- and chemosensory systems, using transcriptomes generated from individual *A. queenslandica* larvae. We first discuss the genes contributing to photo- and chemosensory systems separately; then, we hypothesise how these two sensory systems may be hierarchally integrated at the molecular level to regulate the pelago-benthic transition in this sponge.

### The cryptochrome AqCry2 is part of the photosensory system that prevents settlement when *A. queenslandica* larvae are maintained in the light

Larvae of *A. queenslandica* do not respond to biochemical cues that induce settlement unless they first transition into the dark. Our molecular data indicate a central role for the *cryptochrome (AqCry2*) in this first step of the settlement process, as *AqCry2* showed the largest change in expression coinciding with a light-to-dark transition (Fig. 4*A*). Cryptochromes are blue-light sensing flavoproteins (53) that have a well-characterised role in regulating circadian rhythms in plants (54), insects (55) and mammals (56). The apparent important role for a cryptochrome in light detection and response presented here is consistent with previous findings in this sponge (12) and in other marine invertebrates (45, 47–49, 57). In *A. queenslandica, AqCry2* is spatially expressed in the photo-responsive posterior pigment ring (15) that possesses long cilia that act as phototactic rudders, steering larvae away from light (14). In fact *AqCry2* spatial expression is highest in the pigment ring in “late-ring” larvae that have not yet emerged from the maternal brood chamber (15), supporting their role in the larval phototaxis that is displayed immediately post-emergence (14). Interestingly, after emergence from the dark maternal brood chamber into daylight, *AqCry2* becomes more widely expressed throughout the larva, including in both the outer- and the sub-epithelial layers (15); this is consistent with a far-reaching role during larval life that extends beyond directional swimming via the ciliary rudder. Combined, these data highlight a fundamental role for *AqCry2* in larval photo-responses and are consistent with an overarching role in orchestrating larval settlement.

### The photosensory system prevents settlement by inactivating chemotransduction in *A. queenslandica* larvae, likely via two negative-regulators of G-protein signalling

In *A. queenslandica*, constant light perturbs settlement in response to biochemical settlement cues and retards transcriptional changes through larval ontogeny. Here we discuss three lines of molecular evidence from this study to support the importance of G-protein coupled receptor (GPCR) signalling in the chemosensory system used for sensing and responding to biochemical settlement cues in *A. queenslandica* larvae. First, larvae become responsive to settlement cues after transitioning into the dark (Fig. 2 *A*–*C*), and this is accompanied by a significant decrease in the expression of two putative negative-regulators of G-protein signalling, a GoLoco-motif-containing gene (*AqGoLoco*, Fig. 4*B*) and a regulator of G-protein signalling gene (*AqRGS-GGL*, Fig. 4*C*). Second, 70% *Rhodopsin* and 87% *Glutamate GPCRs* are constitutively expressed throughout larval development indicating a potential role during larval exploration to find a suitable benthic habitat (Fig. S5). Of the GPCRs that were differentially expressed over time, the majority were more highly expressed at 1 hpe compared to 9 hpe (Fig. S6 *A* and *B*), suggesting that *A. queenslandica* larvae may be ‘primed’ to detect biochemical settlement cues upon emergence, but are unable to transduce these cues into a morphogenetic response. Third, of the 18 *Rhodopsin* and 12 *Glutamate GPCRs* that do show significant changes in expression as larvae age, 15 and 9, respectively, only do so only after transitioning into darkness, highlighting a direct connection between light and the expression of these GPCRs (Fig. S6). In fact, two *Rhodopsin GPCRs* are also included in the 180 genes (representing 0.04% of the sponge genome) that are most strongly influenced by light (Fig. 2). Together, these three lines of evidence indicate that *A. queenslandica* larvae detect biochemical settlement cues using GPCRs, but larvae cannot transduce these into a morphogenetic response – which phenotypically manifests as settlement and metamorphosis – until they detect darkness.

GPCRs have previously been hypothesised to regulate molecular pathways underlying larval settlement in this sponge (16) and in other diverse marine invertebrates, including the mollusc *Haliotis rufescens* (6), echinoderm *Stronglylocentrotus purpuratus* (19), arthropod *Balanus amphitrite* (20) and cnidarian *Hydractinia* (21) and *Acropora millepora* (58); but not in the cnidarians *Pocillopora damicornis* and *Montipora capitata* (59) or polychaete *Hydroides elegans* (60). Ligand-bound GPCRs initiate multiple downstream signalling pathways via G-proteins. The three G-protein sub-units (α, β and γ) are disassembled into two different signalling components, an α subunit and a β-γ complex (61). Here, we demonstrate that constant light retards the down- regulation of two genes, *AqRGS-GoLoco* and *AqRGS-GGL*, that are predicted to inactivate the α subunit, indicating strong interdependencies between photo- and chemosensory systems in *A. queenslandica* larvae. We hypothesise that these two inactivators of Gα signalling may maintain larvae in a refractory state, unable to respond to biochemical settlement cues, across space and time until larvae transition into the dark, a physical cue that indicates an appropriate time and place for *A. queenslandica* larvae to settle (Fig. 4*D*).

### The photo- and chemosensory systems may be hierarchically integrated into larval ontogeny via the NO-cGMP pathway

Chemosensory GPCRs are hypothesised to trigger *A. queenslandica* larval settlement via activation of nitric oxide (NO) signalling through the MAPK/ERK pathway (16). Stimulation of NOS produces endogenous NO (62), which either positively or negatively regulates larval settlement in diverse species representing multiple animal phyla (for example, see 16, 63). Indeed, we find that *NOS* is expressed in the top 90th percentile of larval transcript abundance, irrespective of light regime, consistent with its known positive regulation of larval settlement and metamorphosis in this sponge (16). The primary receptor of NO, a soluble guanylyl cyclase (sGC), initiates cyclic guanosine monophosphate (cGMP) signalling cascades (64) and cGMP is also catalysed by the membrane-bound atrial natriuretic peptide (ANP)-activated particulate guanylyl cyclase (ANPR-A) (65). In fact, expression of ANP receptors in the coral *Acropora millepora* indicate an involvement in larval exploration of coralline alga, but not during the morphogenetic responses downstream of an inductive cue (66). In *A. queenslandica* larvae, light perturbs the down-regulation of *ANPR-A* from 1 to 9 hpe and *ANPR-A* is one of the 180 genes (representing 0.04% of the sponge genome) that are most strongly influenced by light. The expression of *ANPR-A* and *NOS* thus support an important role for both NO- and ANP-initiated cGMP signalling cascades for regulating the pelago-benthic transition in *A. queenslandica*. This transition is orchestrated by photo- and chemosensory systems in this sponge, and we propose that both light and biochemical cues may be transduced intracellularly via cGMP signalling cascades that are initiated by *ANPR-A* and *sGC*, respectively. This is consistent with two predicted signal transduction mechanisms, one in mammals and the other in a sponge, that both link photo- and chemosensory systems via NO-cGMP signalling. In mammals, light input via NMDA glutamate receptor (NMDA-R) triggers activation of the NO-cGMP signalling pathway regulating circadian entrainment, and this can intersect with GPCRs via calcium signalling as well (67). In the sponge *Suberites domuncula*, larval photoresponses are hypothesised to be regulated by the photosensory *cryptochrome* that is coupled to the nitric oxide synthase (NOS) pathway, via NOS-interacting protein (NOSIP), and G-protein signalling (68). These two examples thus highlight NO-cGMP signalling as one potential mechanism to explain how photo- and chemosensory systems may be hierarchically integrated at the molecular level to orchestrate the pelago-benthic transition in this sponge.

The current model of NO signalling in *A. queenslandica* involves the GPCR-initiated release of Ca^2+^ from the endoplasmic reticulum into the cytosol in flask cells that are embedded in the outer epithelial-like layer of the larva (16). This Ca^2+^signalling likely allows activation of NOS via binding of calmodulin (64). Here, we provide molecular evidence consistent with that hypothesis, namely that genes coding for proteins involved in Ca^2+^ signalling and Ca^2+^-dependent cellular functions are expressed in a light-dependent manner. Specifically, we find that a sarcoplasmic/ endoplasmic reticulum Ca^2+^ ATPase (SERCA), a plasma membrane Ca^2+^ ATPase (PMCA), and other genes regulating Ca^2+^-dependent cellular functions, are significantly down-regulated over ontogeny, except in the light. These molecular data provide further support that larval responses to light and biochemical cues are interdependent, and may mediate a morphogenetic response through the NO-cGMP signal transduction pathway in this sponge.

### Conclusions

The transcriptome and behavioural data presented here suggest that *Amphimedon queenslandica* larvae may be ‘primed’ to detect biochemical settlement cues upon emergence; indicating that newly emerged *A. queenslandica* larvae may be fundamentally competent to respond to biochemical cues (Fig. S6 *A* and *B*). However, we find that constant light prevents larval settlement, demonstrating deep interdependencies between photo- and chemosensory systems. Our transcriptome data also provide the first molecular evidence to support a recent hypothesis that active inhibition of inducer-signal transduction prevents settlement until such time as larvae detect a physical cue from the environment (8). We hypothesise that the photosensory cryptochrome and chemosensory GPCRs may both converge on the NO-cGMP pathway to regulate the pelago-benthic transition in *A. queenslandica*, via mechanisms as yet undiscovered. The results presented here, for this sponge, phylogenetically extend the importance of interdependent physical and biochemical cues for the pelago-benthic transition far beyond the echinoderms (7, 8) to one of the earliest-branching phyletic animal lineages. In doing so, this study raises the possibility that the requirement of multiple, interdependent sensory systems for regulating larval settlement can be traced back to the last common animal ancestor.

## Methods

### Field collections

For larval behaviour and gene expression assays, adult *Amphimedon queenslandica* sponges were collected from the reef flat of Shark Bay, Heron Island, southern Great Barrier Reef, as previously described (69). Adults were immediately transferred to outdoor aquaria at the Heron Island Research Station (HIRS) supplied with flow-through unfiltered seawater. Sponges were thus subjected to ambient temperature fluctuations (~25-29°C), to biotic and abiotic features of the water pumped into the aquaria from the reef flat, and to a natural day-night cycle; however, aquaria were covered with a light mesh to avoid full sunlight as adults are usually located within crevices and on the undersides of boulders on the reef.

The articulated coralline algae (ACA) *Amphiroa fragilissima* was used as a biochemical settlement cue as this is a known effective inducer of larval settlement in this sponge (10). Clumps of ACA were collected from Heron Island Reef flat and maintained in outdoor flow-through aquaria at HIRS. Pieces of ACA were extracted from multiple different clumps, were lightly brushed to remove debris and epiphytes, and were maintained in 0.22 μm filtered sea water (FSW) overnight, pending experimentation the following day.

### Larval vertical position in the water column through time

To capture larvae that naturally emerge from maternal adults in the early to mid afternoon (11, 12), 16 adult sponges were placed in individual aquaria with larval capture traps fitted to single output hoses. Larvae from different adults were pooled, and 10 individuals randomly collected from the pool were introduced into the top of each 30 ml column (Sarstedt 107×25mm) filled with 0.22 μm FSW via Pasteur pipette. The columns were swirled to disperse larvae and remove any potential bias associated with larval placement in the initial still water conditions. Each assay (described below) was conducted on a different day in February (summer) 2015. To ensure that the counting process did not disturb larval swimming behaviour during hours of ambient darkness, a green filter (LEE filter model 735 velvet green) was used over the light source because *Amphimedon queenslandica* larvae are unresponsive to this wavelength (70). The vertical migration data were analysed using repeated measures ANOVA in SPSS Statistics software v25. Assumptions of normality were satisfied based on graphical analysis of the histograms of the residuals for each time point, except for two time points that had <5 data points. Mauchly’s test indicated that the assumption of sphericity had been violated therefore degrees of freedom were corrected using Greenhouse-Geisser estimates of sphericity (ε) enabling a more conservative F-ratio. These data were visualised using GraphPad Prism version 7.00 for Mac, GraphPad Software, La Jolla California USA, http://www.graphpad.com.

#### Phototaxis on larval vertical migration over time

To assess the effect of dark-shading on the vertical position of *A. queenslandica* larvae in the water column over time, a total of 150, 120 and 60 larvae were collected as described above during three one-hour collection windows beginning at 12, 1 and 2 pm, respectively. These larvae were distributed equally into three different still water experimental conditions within 30 minutes post-emergence. Dark filter paper (LEE filter model 299, 1.2 neutral density) was used to stimulate low light at either the (i) top, (ii) middle or (iii) bottom third of the water with 11 replicate columns for each; these shaded areas were 3 cm in height (~1/3 of the height of the water column). When shading was located at the top and bottom position, the entire top and bottom surfaces of the column were shaded, respectively. The numbers of larvae in the shaded region of each column were recorded hourly for 9 hours, starting at 2 pm, 30 minutes after larvae from the first collection period had been introduced to the into the columns. To determine if shading influenced larval swimming direction over time we used a two-factor repeated-measures ANOVA with time being the within-factor variable and shading position as the between-factor variable.

#### The interaction between photo- and chemo- taxis on larval vertical migration

To determine the effect of light and a biochemical settlement cue on larval vertical migration behaviour through time, a total of 120 larvae were collected from adult sponges from 11:30 am - 12:30 pm and were defined here as being 0 hours post emergence (hpe). Upon emergence, larvae were maintained in plankton mesh reservoirs clipped to the side of a tank, with flowing sea water to mimic natural environmental and temperature fluctuations (~25-29°C). Larvae were distributed equally into the experimental water columns at 4 hpe so as to examine larval positioning in the water column specifically around twilight, in the presence or absence of the ACA covering ~50% of the bottom surface. Larval responses to the presence or absence of ACA were explored under natural day-night and constant light regimes, with each light regime in triplicate, to determine whether the swimming behaviour observed in natural day-night could be altered by constant light (full spectrum Lumilux T5 HE tube). The number of larvae present in the bottom of each column were counted immediately at the start of the experiment (at 4 hpe) and again 30 minutes later (4.5 hpe). Larvae were then counted hourly from 4.5 hpe until 7.5 hpe (~2 hours after sunset) before the final count at 10.5 hpe (11:30 pm). To determine if larval downward migration was affected by light regime and/or cue, these data were analysed using three-factor repeated-measures ANOVA with time being the within-factor variable; cue and light were the between-factor variables.

### Larval settlement under different light regimes

Larvae were collected using a mouth pipette upon their emergence from a pool of 14 adult sponges that were heat-induced (1°C above ambient water temperature) to maximize the numbers of emerging larvae on a single day, in early March (autumn) 2015. To ensure that all larvae were a similar age for gene expression analyses (see below), larvae were collected within a 30-minute window beginning at 11 am and are defined here as being 0 hours post emergence (hpe). To test the effect of different experimental light regimes on larval settlement, 324 larvae were distributed equally into one of three light regimes in either the presence of the ACA or the 0.22 μm filtered sea water (FSW) negative control by 0.5 hpe (n = 54). To ensure that no behavioural interactions or conspecific cues could affect larval settlement responses, each larva was kept in a separate well of 6-well 35-mm diameter sterile polycarbonate tissue culture plates with 10 ml of 0.22 μm FSW either in the presence or absence of two pieces (~5 mm) of the ACA. All settlement plates were semi-submerged in flowing water from the reef, to mimic natural temperature fluctuations, under one of three different light regimes.

The light regimes were (1) natural day-night, (2) constant dark (using LEE filter model 299, 1.2 neutral density, allowing 6.6% transmission of ambient daylight), and (3) constant light (full spectrum Lumilux T5 HE tube). In the constant light regime, settlement assays were conducted on a white surface. Settlement assays under natural day-night were placed on a dark surface to mimic a dark benthic environment (using LEE filter model 299). A small subset of larvae (n = 19) were also transferred from constant light into darkness at 10 hpe, to determine whether settlement behaviour observed in the constant light regime could be altered by transitioning larvae into the dark. Light intensity was measured in triplicate with a Li-Cor LI-1400 data logger that was calibrated with a Li-Cor LI-190 Quantum sensor (Li-Cor, Lincoln, Nebraska, USA).

The numbers of swimming larvae were counted hourly from 1 hpe until 13 hpe, corresponding to 0.5 and 12.5 hours from the time of distribution into their respective light regimes in the presence or absence of a settlement inducer. Any larvae not swimming at the time of counting had settled, which is defined here as attached to the substrate and not dislodged by agitation (swirling) of the plate. To ensure that the counting process did not disturb larval settlement during hours of natural darkness, a green filter (LEE filter model 735 velvet green) was used over a light source because *A. queenslandica* larvae are unresponsive to this wavelength (70).

The effect of light regime and time on settlement was examined with binomial generalised linear models (GLM) defining the binomial response variable as counts of settled and swimming larvae using the R package lme4 (71) and was modelled using a negative binomial distribution allowing for overdispersion. Tukey post-hoc tests (package multcomp; 72) were subsequently performed to conduct pairwise comparisons of larval settlement responses among all light regimes. As no larvae settled in the FSW controls and only 2% had settled in constant light, we used a nonparametric logrank test in the survival package (73) to identify overall significant differences in settlement responses to ACA compared to the FSW controls and to compare settlement when larvae were transferred from the light into darkness at 10 hpe to larvae that remained in the constant light regime.

### The effect of age and light on larval transcriptomes

To test the effect of different light regimes on gene expression, larvae (n=10) were collected every two hours, starting at 1 hpe and ending at 9 hpe, from the FSW negative controls in each light regime that were used in the settlement assays described above. Larvae were immediately rinsed in a 50:50 sea water:RNAlater wash, and fixed in 100% RNA later (Invitrogen) at 4°C for 24 hours to allow the solution to thoroughly penetrate the tissue, before being transferred to −80°C until processing as per manufacturer’s recommendations.

For RNA-sequencing, two timepoints were chosen to represent precompetent and competent larvae, where competency was operationally defined based on low and high rates of settlement (<5% and <50% swimming, respectively) in response to a biochemical inducer in natural day-night; we selected larval ages of 1 and 9 hpe, respectively. To measure gene expression, RNA was isolated from individual larvae using TRIzol (Sigma) (74). RNA quantity and quality were assessed using the Qubit® RNA assay kit on the Qubit® fluorometer (Invitrogen-Life Technologies) and mRNA Pico Series II assay on the Agilent 2100 Bioanalyzer (Agilent Technologies), respectively. The total RNA was stored at −80°C in UltraPure™ DNase/RNase-free water (ThermoFisher) until further processing. From each light regime, we selected 7 larvae with the highest quality RNA (RIN of 8-9) for sequencing, except 8 larvae were sequenced from the natural day-night regime at 9 hpe, because there was one additional space available in the sequencing run. The sample sizes used in this RNA-sequencing experiment were guided by recommendations in (75).

PolyA libraries were prepared according to the CEL-Seq2 protocol (25), using 5 ng starting RNA, and were sequenced on Illumina HiSeq2500 on rapid mode using HiSeq Rapid SBS v2 reagents (Illumina) at the Ramaciotti Centre for Genomics (University of New South Wales, NSW, Australia). The total of 43 larvae were sequenced in a single run on two lanes at a concentration of 8pM. Raw CEL-Seq reads were processed using a publicly available pipeline (https://github.com/yanailab/CEL-Seq-pipeline). The demultiplexed reads were mapped to the *A. queenslandica* Aqu2.1 gene models (28) and read counts were generated. Independent filtering was performed (76) to only include gene models and long non-coding RNAs (lncRNAs) with at least 1 count in 7 or more samples (size of the smallest replicate group).

#### Unsupervised gene expression analyses investigate the relationship between larval age and light on gene expression

To first explore the sources of variation in these gene expression data we used principal component analyses (PCAs) (26) on variance stabilising transformed (vst) counts that were generated in the bioconductor package DESeq2 (27) in R version 3.2.3 (77). This unsupervised analysis does not take into account any information on the sample groups. For this reason, it is a useful tool to understand the sources of variation in gene expression data. The results of this PCA were related to biological information using colour coding with 95% confidence ellipses using ggplot2 in R (78).

Significant changes in gene expression between any level of a factor (light regime and/or time) and their interaction were identified using the LRT analysis in DESeq2 (27). However, the LRT analysis does not produce any metric equivalent to the fold change (FC) that indicates the direction and magnitude of change in expression between two groups. Therefore, we were unable to separate up- and down-regulated genes, as recommended by (79); this is because genes with functional links have positively correlated expression levels and are thus up- or downregulated similarly. Therefore, next we identified significant changes in gene expression over time (adjp < 0.05) using the Wald test in DESeq2 (27). This pairwise comparison was conducted independently for each light regime. Each DEG list was then separated according to whether genes showed a higher expression at 1 hpe (negative FC value) or at 9 hpe (positive FC value). These significantly up- and down-regulated genes were then compared across light regimes using Venny 2.0 (30).

#### Supervised gene expression analysis to identify the ontogenetic changes in gene expression that are perturbed by light

To identify the genes that are most strongly influenced by larval age and light regime we adopted the multivariate sparse partial least squares discriminant analysis (sPLS-DA; 31), implemented in the mixOmics package (80) in R version 3.3.1 (77). This is a supervised analysis that uses the sample information (larval age and light regime) to identify a small subset of genes (a molecular signature) that are most effected by larval age and light regime. To ensure the lowest classification error rate and thus the most predictive model, we identified the optimal number of components as well as the number of parameters (gene models and long non-coding RNAs) to be selected on each component. These components thus capture the changes in gene expression that best explain differences among the 6 sample groups (larvae that were aged 1 or 9 hpe and that were reared under one of the three different light regimes). We obtained a reliable estimation of the error rate of the predictive model by simulating 100 test datasets and performing M-fold cross-validation, whereby the total number of individual larval samples were randomly partitioned into 6 subsets that were used to train and correctly evaluate the performance of the predictive model. The optimised number of components, and parameters to be selected on each component were then included into the final sPLS-DA model. The final model was run on the entire dataset, as recommended in (31), to obtain a single list of differentially expressed genes (DEGs) that best explain variance between sample groups.

To visualise the expression of these genes, we created (i) a sample plot including confidence ellipses (level set to 95%) to highlight the strength of the discrimination between sample groups (using the mixOmics package) and (ii) a heatmap using the R pheatmap package and variance stabilising transformed (vst) counts (81). To separate downregulated and upregulated genes, we identified all genes in each of the two major K-means clusters in the aforementioned heatmap, as each cluster corresponded to higher expression at either 1 or 9 hpe, respectively.

#### Gene annotation and Pfam enrichment analysis

Differentially expressed *A. queenslandica* (Aqu2.1) gene models (28) and long non-coding RNAs (29) were annotated using blastp (e-value cutoff = 1e-3) and InterProScans with the default settings, and these were merged for GO term annotation using Blast2GO (82, 83). Gene models were annotated with the Pfam-A database (84, 85) using a Pfam e-value cutoff of 0.001 and a domain specific e-value of 0.001. KEGG annotations were obtained using the online tool BlastKOALA (86). To ascertain the over-represented functional processes, Pfam enrichment analyses were performed on the sPLS-DA gene list (see above) against a background reference (all genes in the genome) using a previously published R script (87).

*Identification of the NO-cGMP pathway components in* Amphimedon queenslandica *larvae* Larval settlement in *A. queenslandica* may be triggered via GPCR activation of nitric oxide (NO) signalling through the MAPK/ERK pathway (16). To investigate this hypothesis, we identified all previously predicted *Rhodopsin* and *Glutamate GPCRs* (40) and visualised their expression in this RNA-sequencing dataset using a heatmap created in the R pheatmap package (81). We also conducted a pathway analysis to identify the KEGG pathway components specifically relating to the NO-cGMP pathway using the KEGG Mapper – Reconstruct Pathway tool (88). All genes that were expressed in this RNA-seq experiment were used as input into this analysis because the expression of these genes indicates a potential role during larval life.

*Developmental gene expression profiles throughout the* Amphimedon queenslandica *life cycle* For genes that were hypothesised to play a central role in larval settlement, transcript abundance was assessed using CEL-Seq data comprising 82 samples from 17 developmental stages spanning from early cleavage to adult (41, 42; NCBI GEO GSE54364). The normalised gene expression (counts per million mapped) were averaged for each of the 17 developmental stages. These data were visualised using GraphPad Prism version 7 for Mac, GraphPad Software, La Jolla California USA, http://www.graphpad.com.

## Acknowledgments

This research was generously supported by Australian Research Council grants DP110104601 and DP170102353 to SM Degnan. We thank Kerry Roper for assistance with CelSeq library preparation, and Laura Grice and Eunice Wong for constructive comments on the manuscript. We also acknowledge the staff and facilities of the Heron Island Research Station for assistance with fieldwork.

## Supplementary Information

**Fig. S1.**
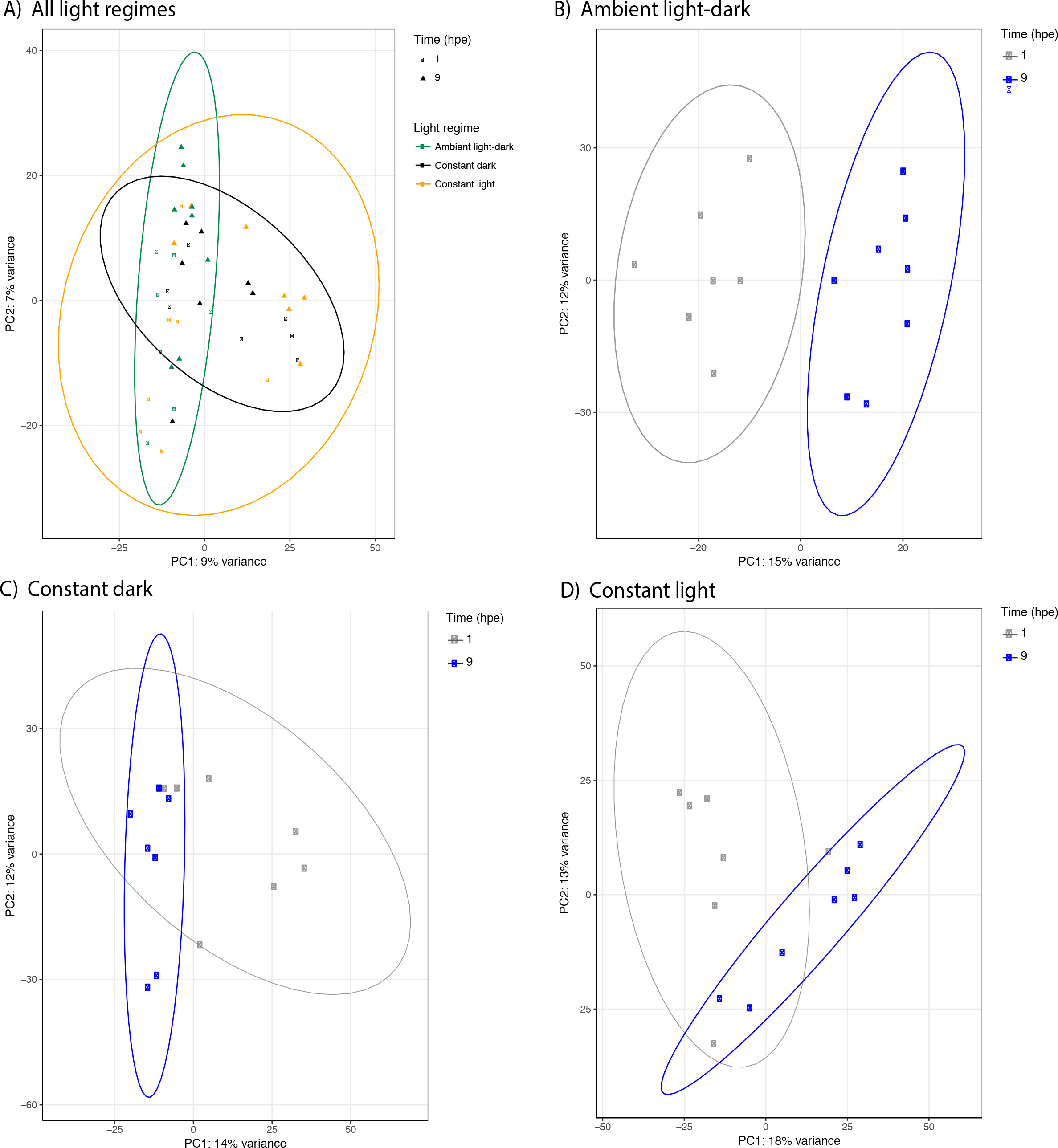
Variation in the transcriptomes of larvae aged 1 and 9 hours post emergence (hpe) that were subjected to three different light regimes. Sources of variation were identified using principal component analyses (PCAs) on variance stabilising transformed (vst) counts and were visualized with 95% confidence ellipses.

**Fig. S2.**
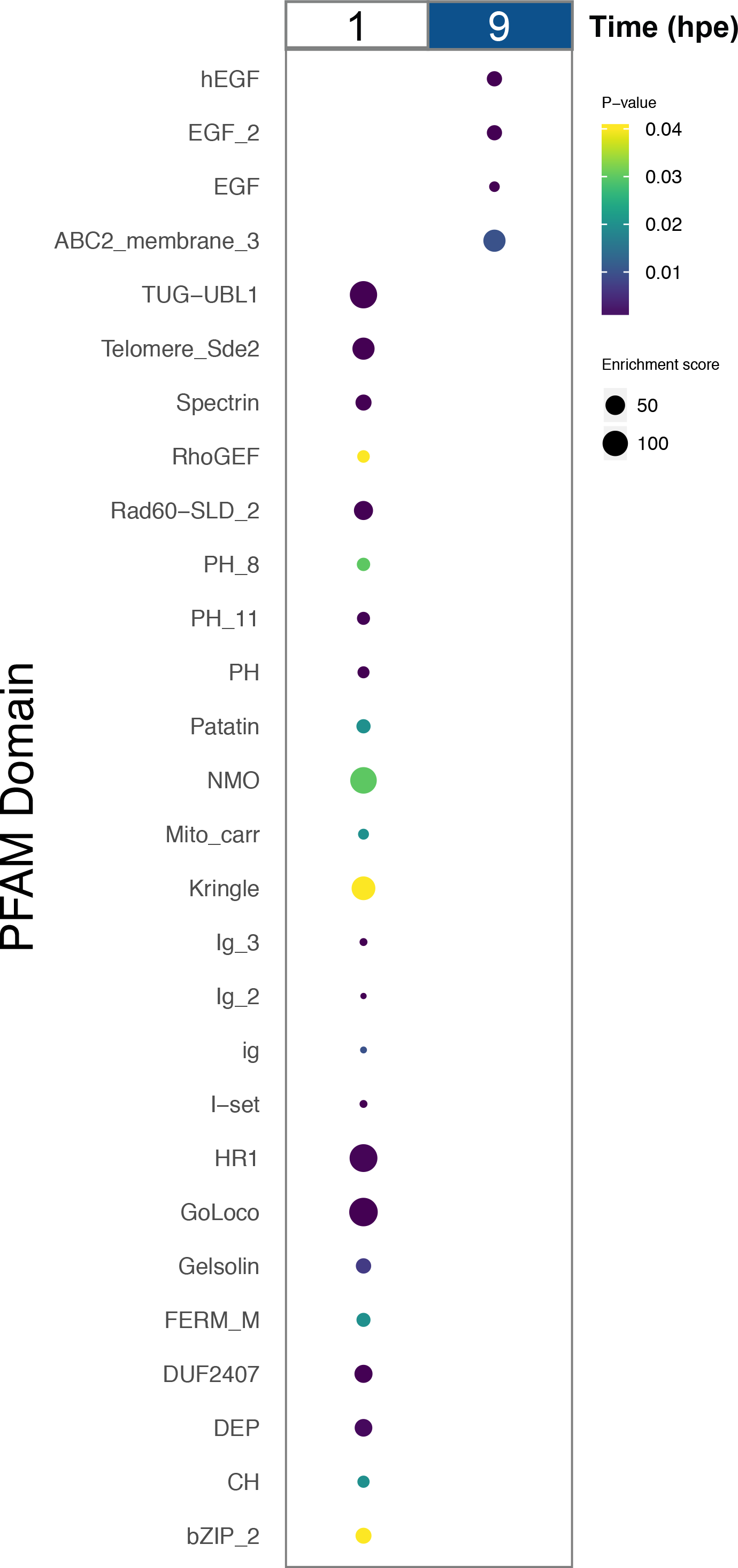
Statistically over-enriched functional Pfam domains in the genes that are most strongly influenced by age and light regime in *Amphimedon queenslandica* larvae. Genes were identified by the sPLS-DA analysis (31) using transcriptomes of larvae aged 1 and 9 hours post emergence (hpe) and reared under constant light, natural day-night, or constant dark. Pfam enrichment analyses were performed against all genes in the genome..

**Fig. S3.**
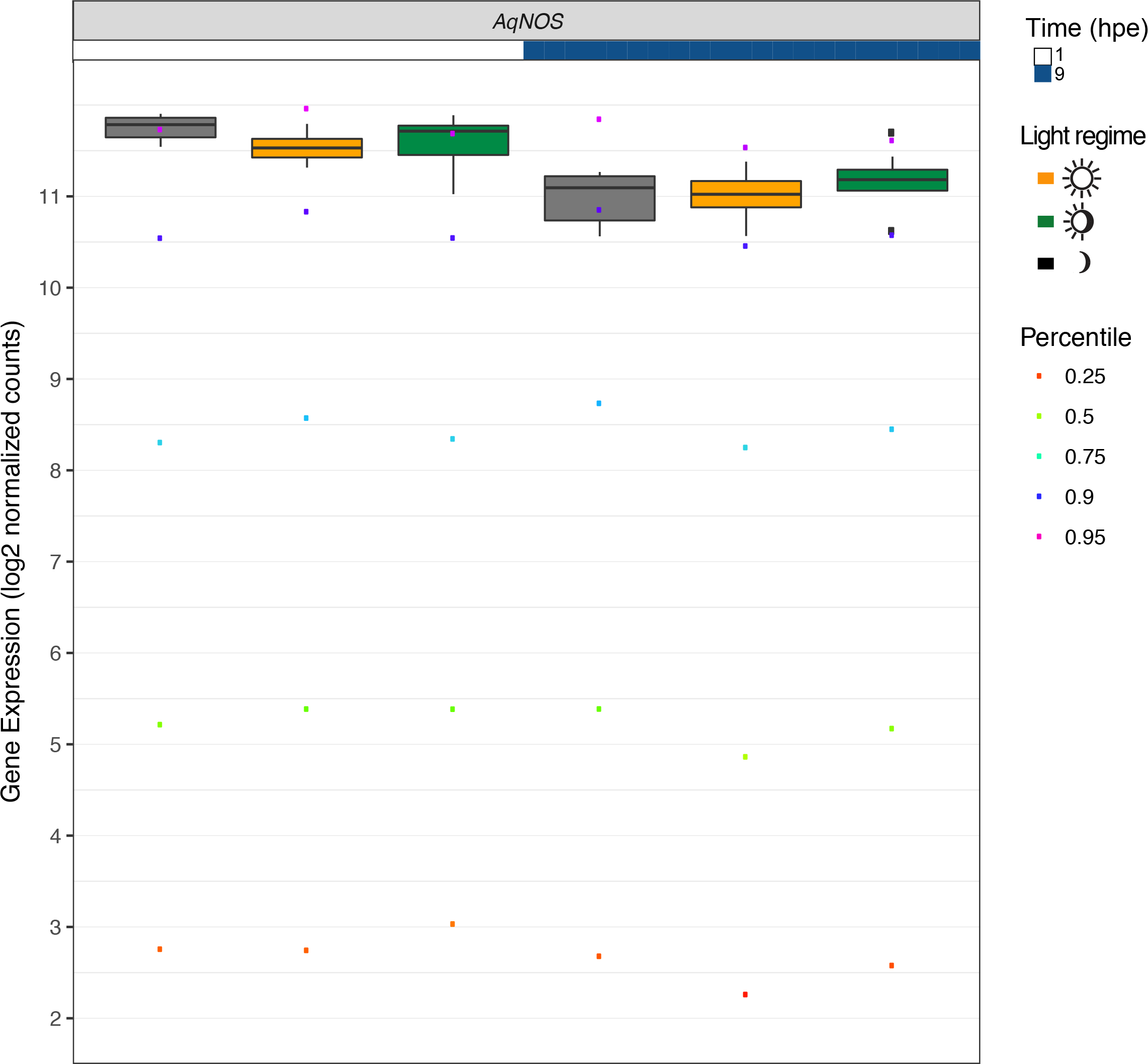
Expression of *AqNOS* in *Amphimedon queenslandica* larvae of two ages that were subjected to three different light regimes relative to transcriptome-wide percentiles. Larvae aged 1 and 9 hours post emergence (hpe) were subjected to constant light, natural day-night, or constant dark. Boxes indicate the interquartile range (IQR) around the median of log_2_ normalised expression and whiskers extend to 1.5 times the IQR. Asterisks represent genes that are significantly differentially expressed over a threshold of 1 (on the log_2_ scale) between 1 and 9 hpe. Coloured dots represent the transcriptome-wide percentiles.

**Fig. S4.**
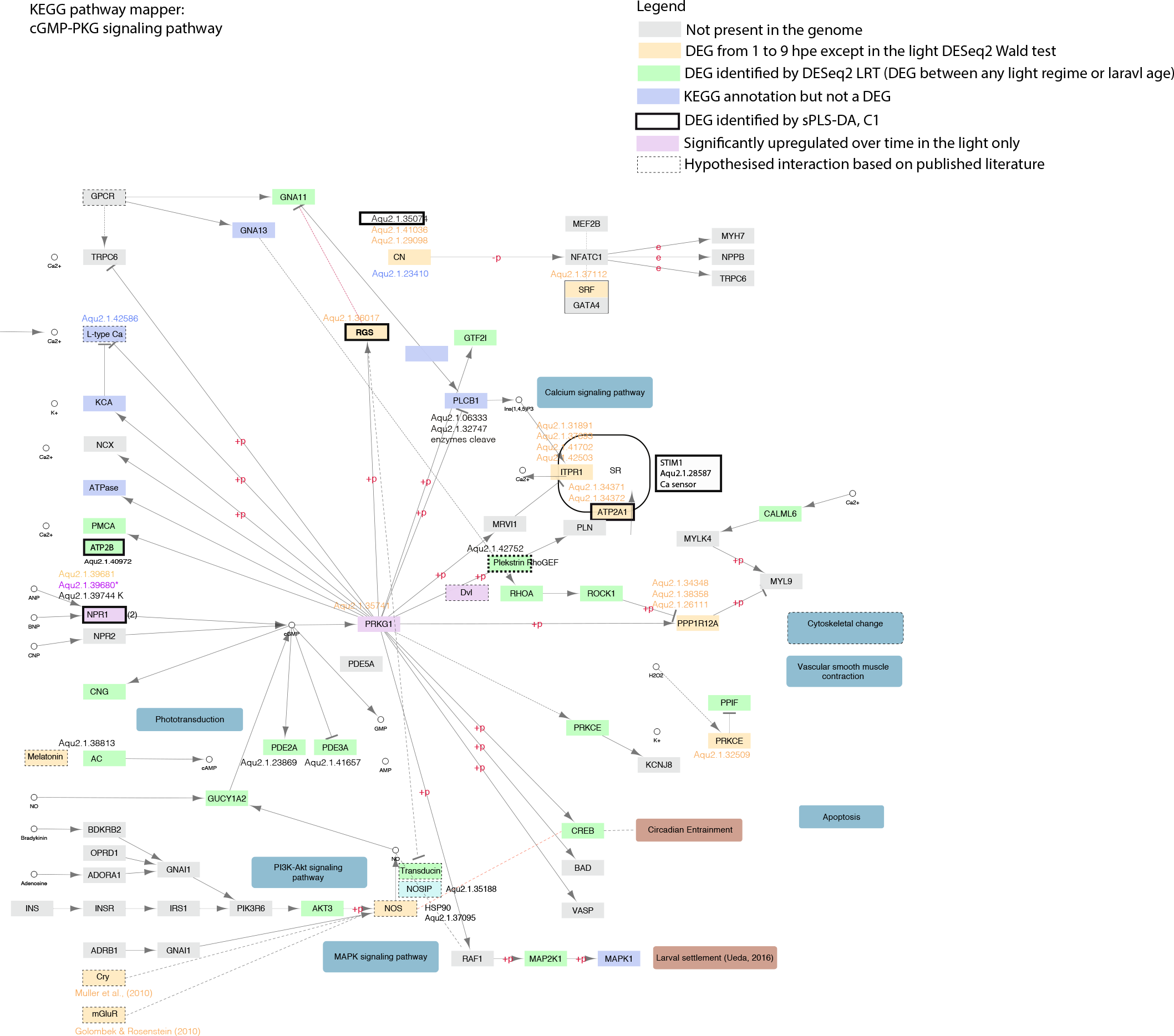
Components of the NO-cGMP signalling human reference pathway from KEGG that are expressed in the sponge *Amphimedon queenslandica*\*. Components of the NO-cGMP signalling human reference pathway that are not present in the *A. queenslandica* genome are highlighted grey. Components that are constitutively expressed in these larval transcriptomic data are highlighted in blue. Differentially expressed genes (DEGs), using a cut off of (adjp < 0.05), are highlighted in orange, green and pink, and each colour denotes a different pairwise comparison. Green denotes DEGs that between any two light regimes, ages and their interaction using the LRT Statistics. Orange denotes DEGs between 1 and 9 hours post emergence (hpe) and pink denotes genes specifically upregulated at 9 hpe in the light, compared to all other light regimes. Thick outline denotes corroboration of the DESeq2 DEGs with the sPLS-DA.

**Fig. S5.**
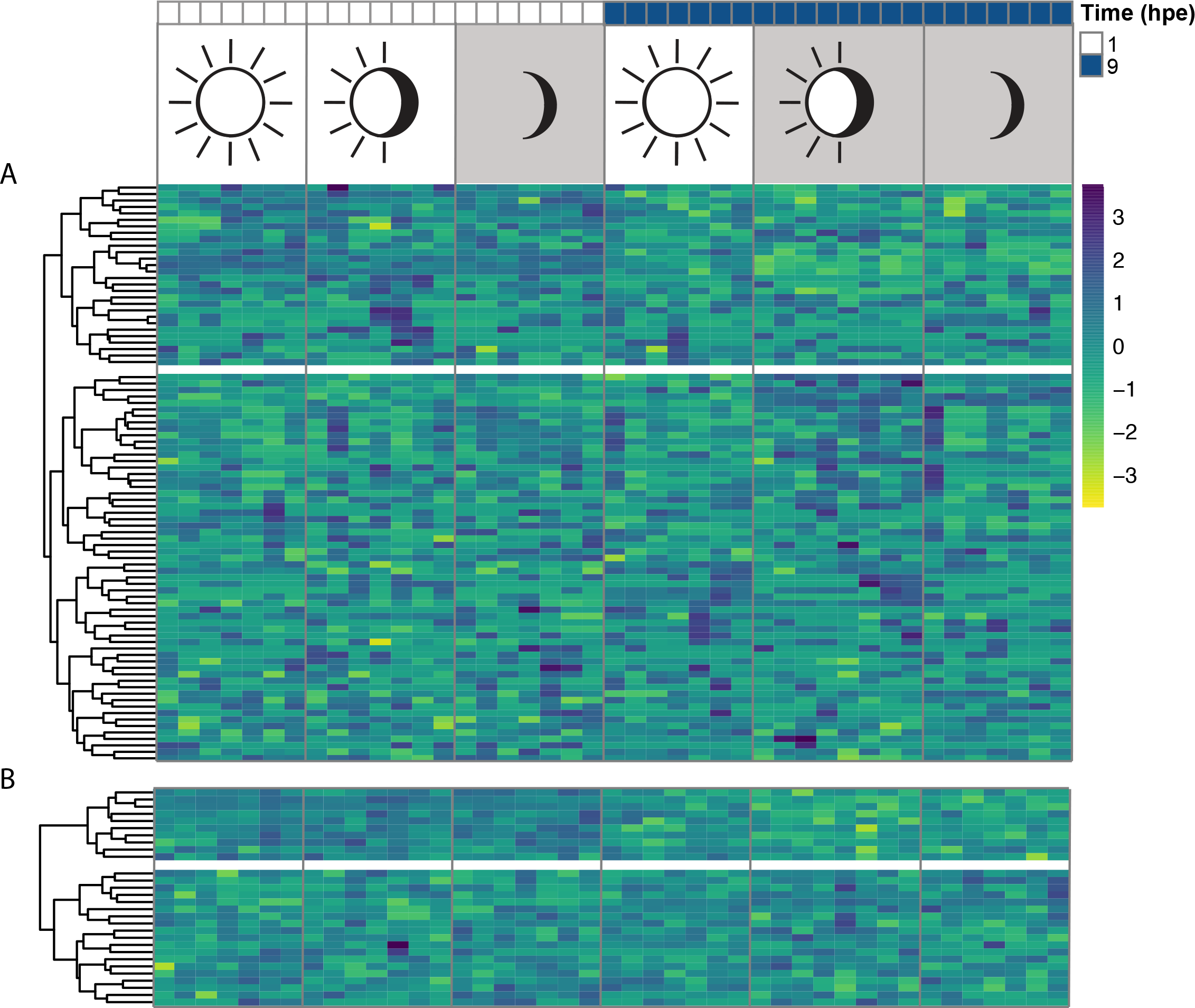
Expression of all (A) *Rhodopsin* and (B) *Glutamate* family of G-protein coupled receptors (GPCRs) in *Amphimedon queenslandica* larvae of two different ages that were subjected to three different light regimes. A hierarchically clustered heatmap of expression using variance stabilising transformed (vst) counts for larvae aged 1 and 9 hours post emergence (hpe) that were subjected to different light regimes (constant light, natural day-night or constant dark).

**Fig. S6.**
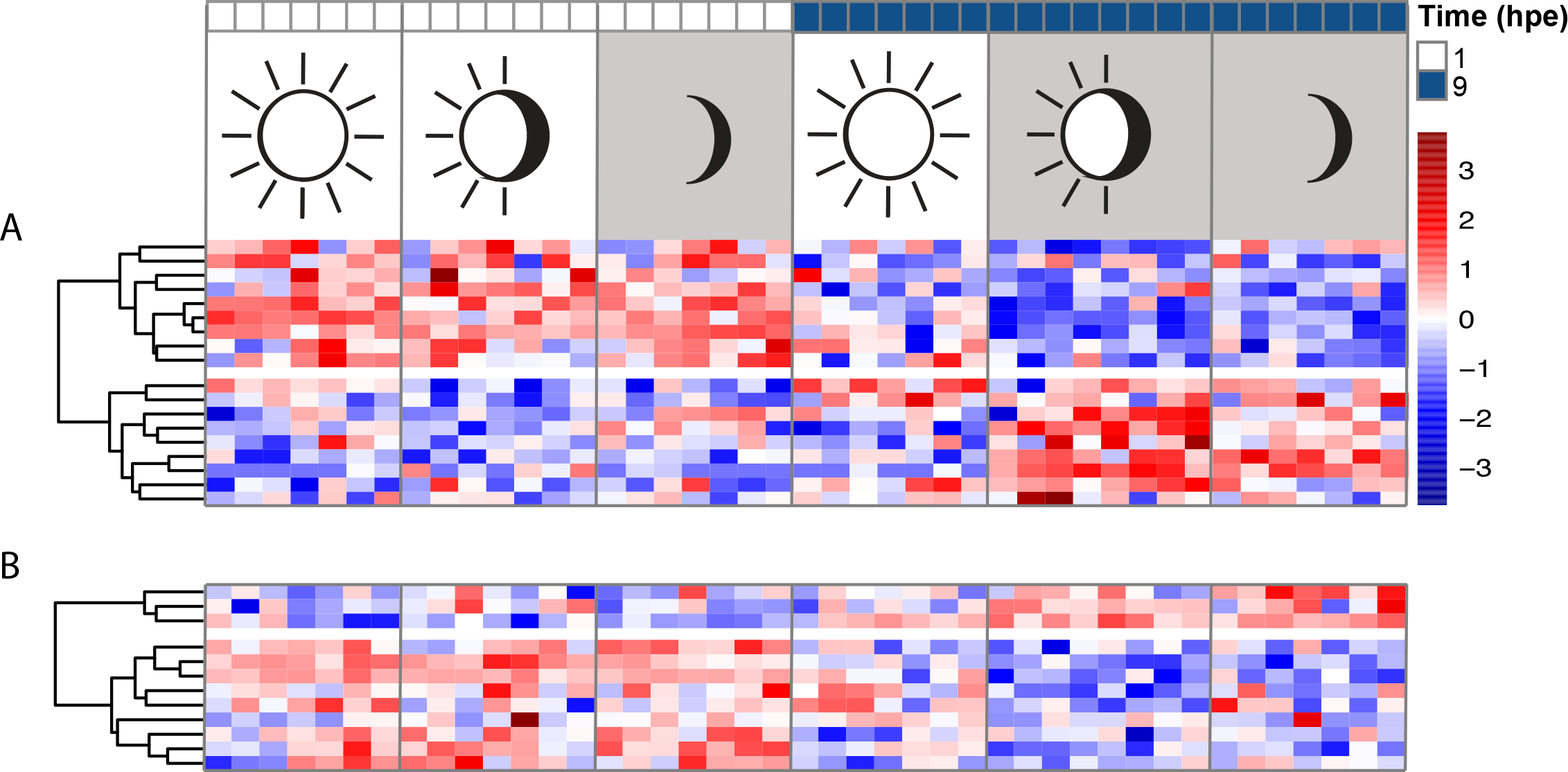
Expression of significantly differentially expressed (A) *Rhodopsin G-protein coupled receptors (GPCRs)* and (B) *Glutamate GPCRs* in larvae of two ages that were subjected to three different light regimes. Larvae were aged 1 and 9 hours post emergence (hpe) and were subjected to different light regimes (constant light, natural day-night and constant dark). The Wald statistical test was conducted in the R package DESeq2 using a cut off of (adjp < 0.05).

**Dataset S1. Summary of the demultiplexing, mapping and counting steps of the CEL-Seq2 pipeline**

**Dataset S2. Raw CEL-Seq reads**

**Dataset S3. Significant changes in larval gene expression between any level of a factor (light regime and time) and their interaction in *Amphimedon queenslandica* larvae**

Significantly differentially expressed genes (DEGs) were identified using the LRT analysis in DESeq2 (27) and were identified at the 0.05 significance level. DEGs were annotated with Blast2GO (82, 83) and KEGG annotations (86). Long non-coding RNAs are annotated as TCONNS.

**Dataset S4. Significant changes in gene expression between larvae aged 1 and 9 hours post emergence (hpe)**

Differentially expressed genes (DEGs) identified using the Wald test in DESeq2 (27) at the 0.05 significance level. This pairwise comparison was conducted for each light regime independently (constant light, natural day-night, and constant dark). Each DEG list was then separated according to whether genes showed a higher expression at 1 hpe (negative FC value) or at 9 hpe (positive FC value). DEGs were annotated with Blast2GO (82, 83), KEGG (86) and Pfam annotations (Pfam-A database; 84, 85). A Pfam e-value cutoff of 0.001 and a domain specific e-value of 0.001 was used. Long non-coding RNAs are annotated as TCONNS.

**Dataset S5. Annotated lists of differentially expressed gene models (DEGs) that show changes in expression through larval ontogeny that are perturbed by light**

The DEGs identified by the sPLS-DA (31) were annotated using Blast2GO (82, 83) and KEGG annotations (86) and were also independently identified by the DESeq2 analysis at the 0.05 significance level (annotated here). Cluster denotes the position in the hierarchically clustered heatmap generated using Pearson correlation on vst transformed counts shown in Fig. 3*B* (1= top, 2= bottom cluster). Long non-coding RNAs are annotated as TCONNS.

**Dataset S6. Pfam annotations for the complete lists of differentially expressed gene models (DEGs) and long non-coding RNAs that show changes in expression through larval ontogeny that are retarded by constant light**

These DEGs were identified by the sPLS-DA (31) and were annotated with the Pfam-A database (84, 85) using a Pfam e-value cutoff of 0.001 and a domain specific e-value of 0.001. Long non-coding RNAs are annotated as TCONNS.

**Dataset S7. Complete lists of *Rhodopsin* and *Glutamate* families of *G-protein coupled receptors (GPCRs)***

These GPCRs that were originally identified by (40) have a match to the Aqu2.1 gene models (28) and are annotated here with the Pfam-A database (84, 85) using a Pfam e-value cutoff of 0.001 and a domain specific e-value of 0.001.

